# The yeast Mig1 transcriptional repressor is dephosphorylated by glucose-dependent and independent mechanisms

**DOI:** 10.1101/130690

**Authors:** Sviatlana Shashkova, Adam J.M. Wollman, Mark C. Leake, Stefan Hohmann

## Abstract

*Saccharomyces cerevisiae* AMPK/Snf1 regulates glucose derepression of genes required for utilization of alternative carbon sources through the transcriptional repressor Mig1. It has been suggested that the Glc7-Reg1 phosphatase dephosphorylates Mig1. Here we report that Mig1 is dephosphorylated by Glc7-Reg1 in an apparently glucose-dependent mechanism but also by a mechanism independent of glucose and Glc7-Reg1. In addition to serine/threonine phosphatases another process including tyrosine phosphorylation seems crucial for Mig1 regulation. Taken together, Mig1 dephosphorylation appears to be controlled in a complex manner, in line with the importance for rapid and sensitive regulation upon altered glucose concentrations in the growth medium.

## Introduction

Budding yeast *Saccharomyces cerevisiae* adapts its metabolism to various environmental stress factors through the AMP-activated protein kinase (AMPK)/Snf1 protein kinase (Carling, et al. 1994). The main role of Snf1, the adaptation of the cell to glucose limitation via the glucose derepression pathway (Carlson, et al. 1981, Celenza and Carlson 1986), is executed through the control of the transcriptional repressor Mig1 (Carlson 1999). Under glucose-rich conditions, Snf1 kinase activity is inhibited mainly through dephosphorylation by the Glc7 phosphatase associated with its regulatory subunit Reg1 (McCartney and Schmidt 2001, Tu and Carlson 1994). Reg2, a paralogue of Reg1, is another Glc7 regulatory subunit which participates in negative regulation of Snf1 (Frederick and Tatchell 1996, Jiang, et al. 2000, Maziarz, et al. 2016). Sit4 and Ptc1 phosphatases have also been suggested to contribute to Snf1 dephosphorylation (Rubenstein, et al. 2008, Ruiz, et al. 2012).

In high glucose conditions Mig1 becomes dephosphorylated and relocates to the nucleus (Treitel and Carlson 1995, Wu and Trumbly 1998) where it occupies promoters of target genes (*SUC2*, *GAL*, *MAL*) resulting in their repression (Hu, et al. 1995, Johnston, et al. 1994, Nehlin, et al. 1991, Wu and Trumbly 1998). Upon glucose limitation, Snf1 becomes activated by phosphorylation on Thr210 (McCartney and Schmidt 2001, Nath, et al. 2003). Active Snf1 in turn phosphorylates Mig1 on multiple sites (DeVit and Johnston 1999, Treitel, et al. 1998) leading to Mig1 dissociation from the DNA and apparent Mig1 relocalization to the cytoplasm (De Vit, et al. 1997, DeVit and Johnston 1999, Elbing, et al. 2006, Hong, et al. 2003, Nath, et al. 2003, Rubenstein, et al. 2008). Although, the Snf1/Mig1 glucose repression pathway has been intensively studied and potential cross talk with other pathways has been reported (Shashkova, et al. 2015), the exact mechanism of Mig1 regulation remains unclear.

NaF and Na_3_VO_4_ phosphatase inhibitors are routinely included in extraction buffers to preserve protein phosphorylation state. Vanadate anions are known to inhibit tyrosine phosphatases (Gordon 1991) due to structural similarity to orthophosphate ions (Crans, et al. 2004, Korbecki, et al. 2012). Na_3_VO_4_ is also an inhibitor of other enzymes including alkaline phosphatases and ATPase. NaF, another protein phosphatases inhibitor, prevents dephosphorylation on serine/threonine residues (Shenolikar and Nairn 1991). It has been shown that NaF does not change the total activity of these phosphatases but selectively inhibits some of them. For instance, inhibition of myosin-specific phosphatase results in endothelial cell barrier function via its effect on actin (Wang, et al. 2001).

While it is well established that Glc7-Reg1/2 dephosphorylates Snf1 it is less clear which phosphatases act on Mig1 and if the Mig1 phosphorylation status is solely determined by Snf1 activity. A possible role for the Glc7-Reg1 phosphatase has been suggested (Rubenstein, et al. 2008). In this study we employed a combination of biochemical assays and cutting-edge single-molecule imaging approaches to investigate a link between the Mig1 phosphorylation status and its cellular localization under different glucose conditions. Mig1 localization, as characterized by standard fluorescence microscopy, has been extensively used as a marker for phosphorylation and signaling activity (Bendrioua, et al. 2014). Here, we use Slimfield microscopy combined with deconvolution software to determine the molecular concentration of Mig1 in the nucleus and cytoplasm in single live cells (Wollman and Leake 2015). We propose two different (glucose-dependent and independent) mechanisms of Mig1 dephosphorylation and both in combination seem to be required for complete Mig1 dephosphorylation.

## Materials and methods

### Growth conditions

Standard YPD (10 g/l yeast extract, 20 g/l bacto-peptone, glucose according to experimental needs) and YNB (1.7 g/l yeast nitrogen base without amino acids, without (NH_4_)_2_SO_4_, 5 g/l (NH_4_)_2_SO_4_, supplemented with glucose and amino acids according to nutritional requirements) were used for yeast cell growth and transformant selection. For all experiments cells were pre-grown overnight in appropriate media at 30^0^C, 180 rpm. *S. cerevisiae* strains and plasmids used in this study are listed in Table 1.

### Strain construction

The *LEU2* fragment from YDp-L plasmid flanked on its 5’- and 3’-ends with 50 bp upstream and downstream of *SNF1*, respectively, was amplified by PCR. Strain YSH2348 was transformed directly with the PCR reaction mix by standard LiAc protocol (Gietz and Schiestl 2007). Prior to placing onto YNB plates without leucine, the transformants were incubated in YPD supplemented with 4% glucose for 1h at 30^0^C, 180 rpm.

### Protein extraction

Yeast cells carrying p*MIG1-HA* and p*SNF1-TAP* or *pSNF1-I132G-TAP* plasmids were grown until mid-logarithmic phase on appropriate media with 4% glucose (w/v). Cultures were then split into two different culture flasks and supplemented with 4% or 0.2% glucose media for 30 min. 25μM 1NM-PP1 (Cayman) or 10mM Na_3_VO_4_ were added for 5 min at room temperature, 0.1M NaOH was then added to the cultures for 5 min. For NaF treatment, prior to NaOH addition, cultures were incubated with 10mM NaF for 2h at 30^0^C while shaking. For the time course experiments with 1NM-PP1, NaOH was added at desired time intervals. Cells were harvested by centrifugation (3000 rpm, 50s), the pellets were suspended in 2M NaOH with 7% β-mercaptoethanol for 2 min and then 50% trichloroacetic acid was added. Samples were vortexed and spun down at 13 000 rpm. The pellets were washed in 1M Tris-HCl (pH 8.0), suspended in 50 μl of 1x SDS sample buffer (62.5mM Tris-HCl, 3% SDS, 10% glycerol, 5% β-mercaptoethanol, and 0.004% bromophenol blue, pH 6.8) and boiled for 5 min. The protein extracts were obtained by centrifuging and collecting the supernatants. 70 μg of total protein extract was incubated with 240 units of Lambda protein phosphatase (λPP) (NEB) with 1mM MnCl_2_ buffer at 30°C for 30 min, the reaction was then stopped by adding 200mM Na_3_VO_4_.

### Western blotting

70 μg of total protein extract was loaded on a Criterion TGX Stain Free precast gel (10% acrylamide, Bio-Rad), separated by SDS-PAGE at 300 V for 25 min and transferred onto a nitrocellulose membrane (Trans-Blot Turbo Transfer Pack, Bio-Rad) using Trans-Blot Turbo Transfer System (Bio-Rad). The membrane was washed and blocked in Odyssey buffer (LI-COR Biosciences) for 1h. Mig1-HA was probed with primary mouse anti-HA antibodies (1:2000, Santa Cruz), then secondary fluorescent goat anti-mouse IRDye-800CW antibodies (1:5000, LI-COR Biosciences) and detected by an infrared imager (Odyssey, LI-COR Biosciences), 800 nm channel.

### Epifluorescence microscopy

Pre-grown cells were cultivated in YNB medium with complete amino acid supplement with 4% or 0.2% glucose for 1 h. 1NM-PP1 was added to the cell cultures as described above. For the wide field fluorescence microscopy cells were imaged using an ApoTome camera and a Zeiss Axiovert 200M microscope (Carl Zeiss MicroImaging). Fluorescence images were acquired by using separate filter sets 38HE and 43HE for GFP and mCherry excitation, respectively.

### Slimfield microscopy and spots detection

Pre-grown cells were cultivated in YNB medium with complete amino acid supplement with 4% or 0.2% glucose for 1 h, then surface immobilized on a 1% agarose pad with desired nutrients and glucose concentrations between a microscopy slide and a plasma clean coverslip (Wollman 2016). NaF, 1NM-PP1 or Na_3_VO_4_ were added to the cultures prior to imaging as described above. Simultaneous Slimfield illumination (Plank, et al. 2009) was obtained using co-aligned 488 nm and 561 nm 50mW lasers (Coherent Obis) de-expanded to direct a beam onto the sample at 6W/cm^2^ excitation intensity.

For fluorescence emission acquisition we used a 1.49 NA oil immersion objective lens (Nikon). The total fluorescence signal was split into green and red detection channels using a dual-pass green/red dichroic mirror centered at long-pass wavelength 560nm and emission filters with 25nm bandwidths centered at 525nm and 594nm (Chroma). Each channel was imaged separately at 5ms exposure time by the EMCCD camera (iXon DV860-BI, Andor Technology, UK) with 80 nm/pixel magnification.

Fluorescent spots, within the microscope depth of field were detected and quantified using a single particle tracking algorithm with ∼40nm precision (Miller, et al. 2015, Wollman and Leake 2015) adapted from similar studies reported previously (Beattie, et al. 2017, Reyes-Lamothe, et al. 2010). The mCherry fluorescence signal under glucose repletion was used as a marker for the nucleus. The intensity of spots was characterized using step-wise photobleaching methods, to determine the intensity of a single GFP-tagged molecule (Wollman, et al. 2016) and compared to the intensity of single GFP and mCherry molecules identified in *in vitro* immobilized protein assays described elsewhere (Leake, et al. 2006). This allowed the copy numbers of fluorescent proteins to be determined by the CoPro convolution algorithm (Wollman and Leake 2015).

## Results

### Mig1 dephosphorylation is regulated by two mechanisms

Under glucose rich conditions Mig1 together with two corepressors, Ssn6 and Tup1, occupies and represses promoters of target genes (Treitel and Carlson 1995). Mig1 phosphorylation mediated by Snf1 upon glucose limitation has been shown to be essential for disassembly of this repressor complex and Mig1 export to the cytoplasm (De Vit, et al. 1997, DeVit and Johnston 1999, Papamichos-Chronakis, et al. 2004). Therefore, we first examined Mig1 localization when Snf1 kinase activity is blocked. We employed cells expressing Snf1-I132G, a PP1 analogue-sensitive version of Snf1, with GFP-tagged Mig1 and Nrd1-mCherry as a reporter for the nucleus. Previous studies have shown that wild type forms of kinases are insensitive to PP1 analog inhibitors (Knight and Shokat 2007, Rubenstein, et al. 2008) but a mutation at the ATP-binding site of a kinase to alanine or glycine creates a pocket with a novel structure sensitive to ATP competitive kinase inhibitor 1NM-PP1 (Knight and Shokat 2007, Shirra, et al. 2008). This analogue-sensitive version of Snf1, Snf1-I132G, is fully functional and has been extensively used and well characterized (Rubenstein, et al. 2008, Shirra, et al. 2008, Zaman, et al. 2009).

Incubation of cells expressing Snf1-I132G with 25 μM 1NM-PP1 for 5 min or longer, resulted in Mig1 nuclear localization even under low glucose conditions (Figure 1A), whereas cells with wild type Snf1 were unresponsive to the inhibitor (Supplementary figure 1 A). This observation confirms that Snf1 kinase activity is essential for Mig1 nuclear export under glucose depletion and that the Snf1-inhibitor pair functions as expected in this context.

**Figure 1.**
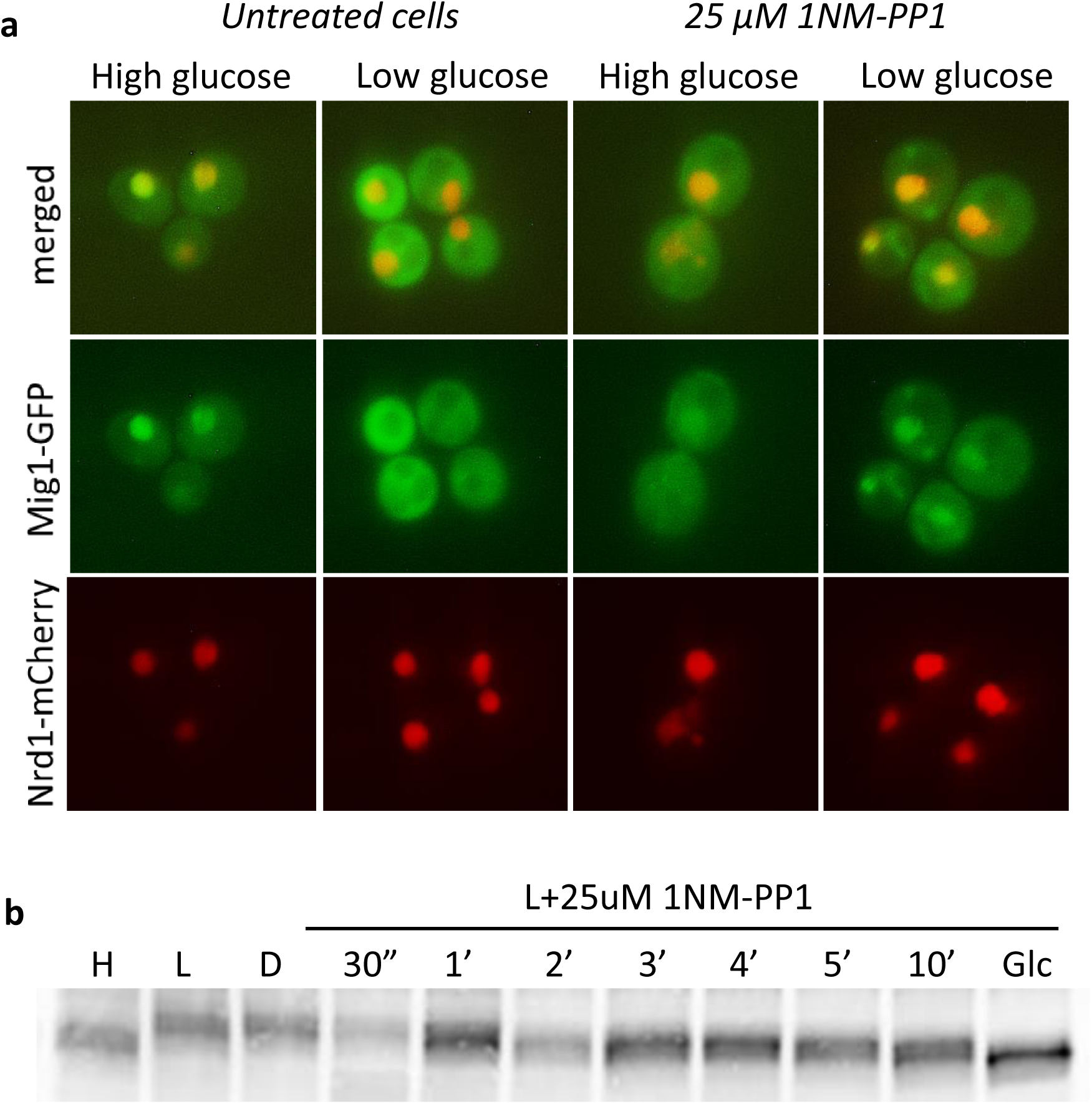
Pharmacological inhibition of Snf1 results in Mig1 dephosphorylation and nuclear localisation upon glucose depletion. **(a)**. Cells carrying genomicaly integrated Mig1-GFP and Nrd1-mCherry fusions and Snf1-I132G we incubated in different glucose conditions with subsequent addition of 1NM-PP1 for 5 min prior to imaging. (b) Cells were grown in 4% glucose, shifted to 4% (H) and 0.2% glucose (L) for 1h. The low glucose cultures were subsequently treated with either DMSO (D) for 5 min, or 25uM 1NM-PP1 (NM) for 0.5, 1, 2, 3, 4, 5, 10 min. 4% glucose (Glc) was6added to a sample treated with 1NM-PP1 for 10 min.

We also checked the effect of Snf1 kinase activity on Mig1 phosphorylation. Mig1 appears as a range of different bands. We assume that those bands correspond to different phosphorylated forms, which is consistent with a theory suggested previously (Needham and Trumbly 2006). We analyzed whether mutated Snf1 affects the phosphorylation status of Mig1 (Supplementary figure 1 B). Our results showed no difference in Mig1 band shift under high and low glucose between the wild type form of Snf1 and Snf1-I132G. A time course experiment indicated gradual dephosphorylation of Mig1 in cells under pharmacological inhibition (Figure 1B). Maximal Mig1 dephosphorylation caused by 1NM-PP1 in low glucose is obtained after 2 min of incubation with the inhibitor, however, several phosphorylated forms of Mig1 can still be observed. Complete Mig1 dephosphorylation was detected only when in addition to Snf1 inhibition, glucose levels were elevated. This suggests that Mig1 dephosphorylation can be mediated by a glucose-regulated mechanism as well as a glucose-independent mechanism.

### Glucose-dependent dephosphorylation of Mig1 is mediated by Reg1

To analyze whether Glc7-Reg1 plays a role in Mig1 dephosphorylation, we performed western blot analysis on strains carrying deletions of *REG1*, its paralogue *REG2* and the *reg1⊿ reg2⊿* double mutant (Figure 2). All strains expressed Snf1-I132G and Mig1-HA. Levels of Mig1 phosphorylation in high (lanes 1, 6, 11 and 16) and low glucose (lanes 2, 7, 12 and 17), respectively, appear to be the same for all strains regardless of Reg1 presence. Similarly, inhibition of Snf1 with 1NM-PP1 resulted in no differences between strains in low glucose conditions (lanes 4, 9, 14 and 19). Interestingly, rapid elevation of the glucose level from 0.2% to 4% resulted in greater apparent dephosphorylation in the wild type and the *reg2⊿* mutant (lanes 5 and 15), compared to initial high glucose conditions (lanes 6 and 16), but not in the *reg1⊿* strain (lanes 10 and 20). Similarly to Figure 1B, total Mig1 dephosphorylation in Reg1 deficient strains was only achieved upon 4% glucose and 1NM-PP1 presence (Supplementary Figure 2). These data suggest that glucose dependent Mig1 dephosphorylation is driven by Glc7-Reg1 phosphatase, but to achieve complete Mig1 dephosphorylation a glucose-independent mechanism is also involved.

**Figure 2.**
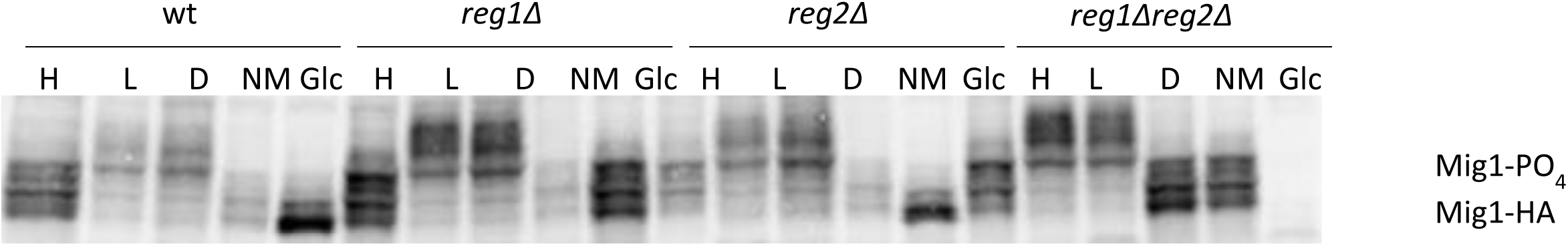
Mig1 dephosphorylation by Glc7-Reg1 is mediated by glucose. The phosphorylation status of Mig1 was determined by Western blotting of protein extracts from cells expressing HA-tagged Mig1 and an ATP-analogue sensitive Snf1, Snf1-I132G, grown in 4% glucose, shifted to 4% (H) and 0.2% glucose (L) for 1h. 0.2% glucose cultures were subsequently treated with DMSO (D), the inhibitor of Snf1-I132G, 25uM 1NM-PP1 (NM), or 4% glucose (Glc) for 5 min.

### Incubation with Na_3_VO_4_ results in Mig1 phosphorylation under high glucose conditions

We also investigated the effects of phosphatases inhibitors on Mig1 phosphorylation. To study their influence, we performed SDS-Page and subsequent western blotting with anti-HA antibodies on total protein extracts containing Snf1-I132G. Mig1 from cells exposed to rapid glucose elevation and with inhibited Snf1 activity was assumed as totally unphosphorylated. 10mM Na_3_VO_4_ or NaF was added to the cultures prior to harvesting cells and protein extraction. Surprisingly, the tyrosine phosphatase inhibitor, Na_3_VO_4,_ showed Mig1 phosphorylation even under glucose repletion (Figure 3A) but the serine/threonine phosphatase inhibitor, NaF, showed no effect (Figure 3B). In control experiments, incubation with λPP resulted in Mig1 dephosphorylation under all conditions.

**Figure. 3.**
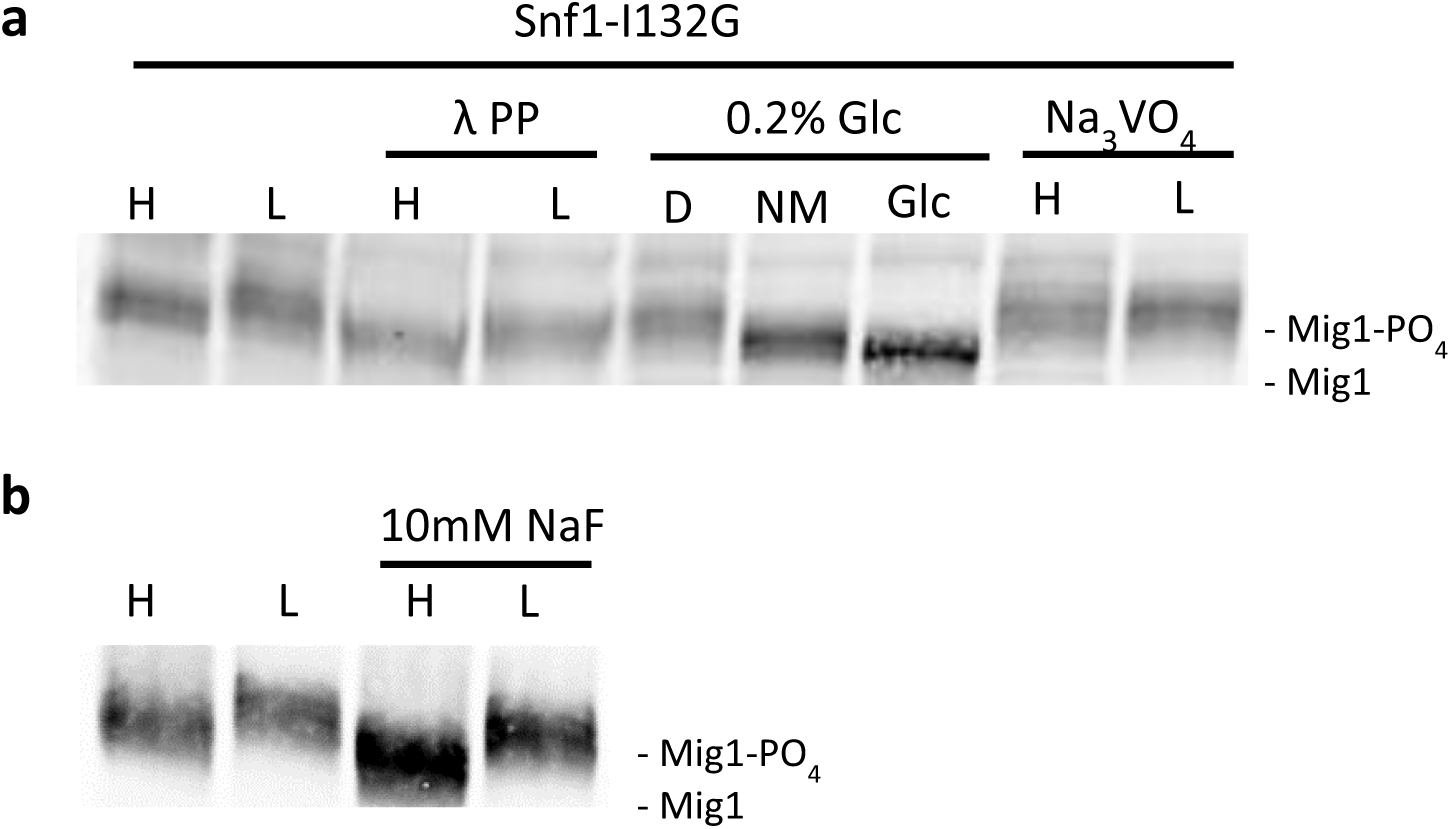
Tyrosine phosphorylation plays a role in Mig1 regulation. Phosphorylation status of Mig1 obtained from 70 μg of total protein extracts from cells expressing Mig1-HA and analog-sensitive version of Snf1, Snf1-I132G, under 4% (H) or 0.2% (L) glucose conditions. **(a, b)** Cultures were exposed to 10mM Na_3_VO_4_, DMSO (D) or 25μM 1NM-PP1 (NM) for 5 min prior to protein extraction. Glucose was elevated to 4% (Glc) in cultures containing 0.2% glucose for 5 min prior to protein extraction. Total cell protein extracts were incubated with Lambda protein phosphatase (λPP) for 30 min at 30 0C. **(c)** Pre-grown cells were incubated with 10mM NaF for 2h prior to protein extraction.

Thus, while we expected that the serine/threonine phosphatase inhibitor would affect Mig1 dephosphorylation under glucose repletion, sodium ortho-vanadate but not fluoride resulted in Mig1 phosphorylation. Although, the effect of NaF might be weak and introduced only on few Ser/Thr sites which would be difficult to capture by western blotting. At the same time, serine/threonine/tyrosine λPP is able to dephosphorylate Mig1. Taken together these data suggest that another factor is responsible for keeping Mig1 unphosphorylated in high glucose in addition to serine/threonine phosphatases. This mechanism of Mig1 dephosphorylation is triggered by a tyrosine specific phosphatase. However, no Tyr phosphorylation sites of Mig1 have been identified.

### Incubation with NaF alters Mig1 cellular localization

We used fluorescence microscopy to test the effects of inhibitor compounds on Mig1 localization but using a single-molecule Slimfield microscope which, combined with deconvolution software, allowed us to measure the absolute Mig1 concentration in the nucleus and cytoplasm. This level of quantitation was necessary to discern any subtle changes to Mig1 localization, undetectable with standard fluorescence microscopy. Strains containing Mig1-GFP and Nrd1-mCherry were imaged in different conditions (Figure 4) and analyzed. The distribution of Mig1 concentrations in the nucleus and cytoplasm across a population of cells was rendered using kernel density estimation (Figure 5).

**Figure. 4.**
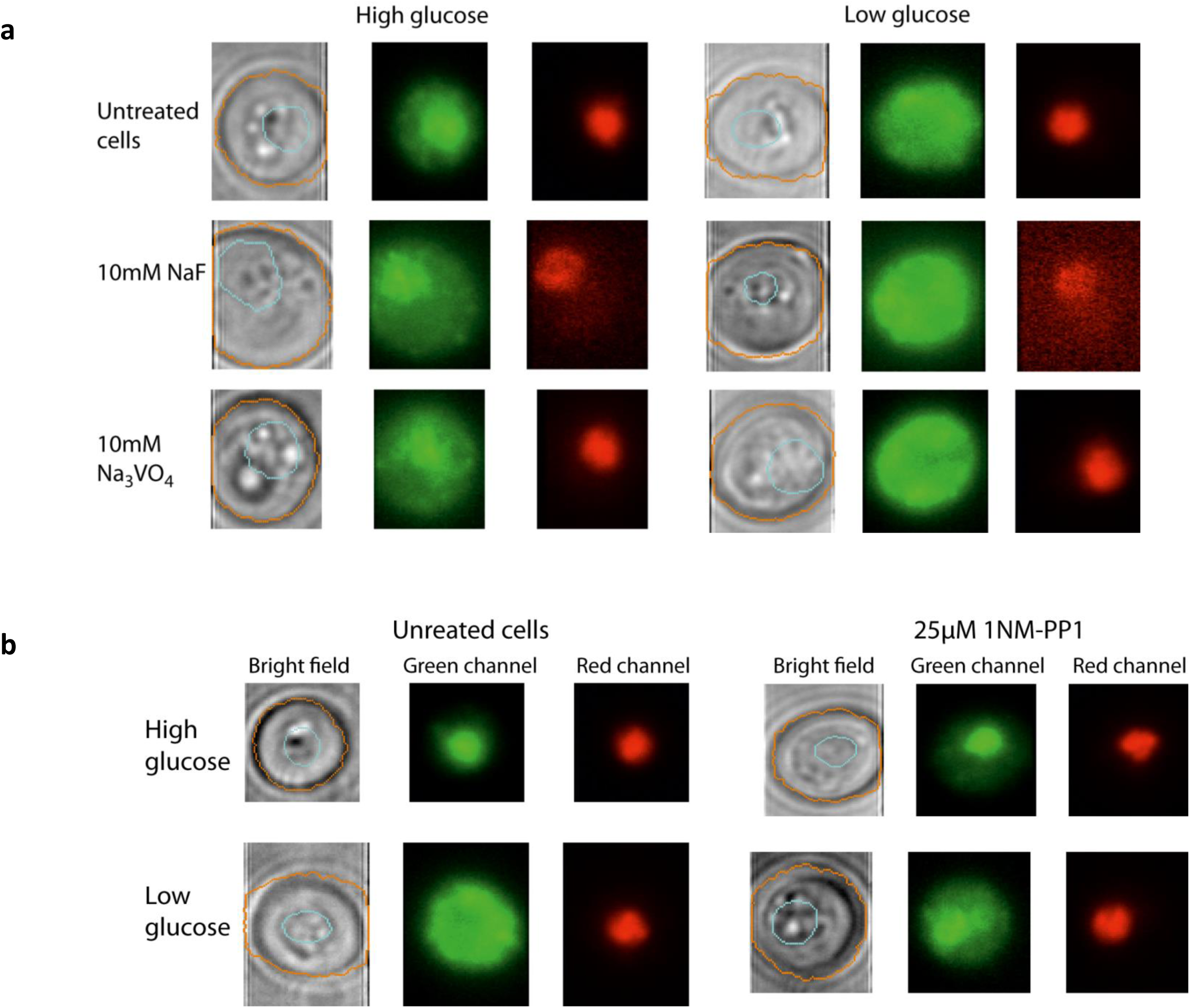
Mig1 localizaion is affected by phosphatases and Snf1. Mig1-GFP localization under 4% (H) or 0.2% (L) glucose conditions. Nrd1-mCherry is a marker for the nucleus location. **(a)** Cultures were exposed to 10mM Na_3_VO_4_, or 10mM NaF for 2h prior to imaging. **(b)** Pre-grown cells carrying Snf1-I132G were incubated in low glucose and then exposed to 25μM 1NM-PP1 for 5 min prior to Slimfield imaging.

**Figure. 5.**
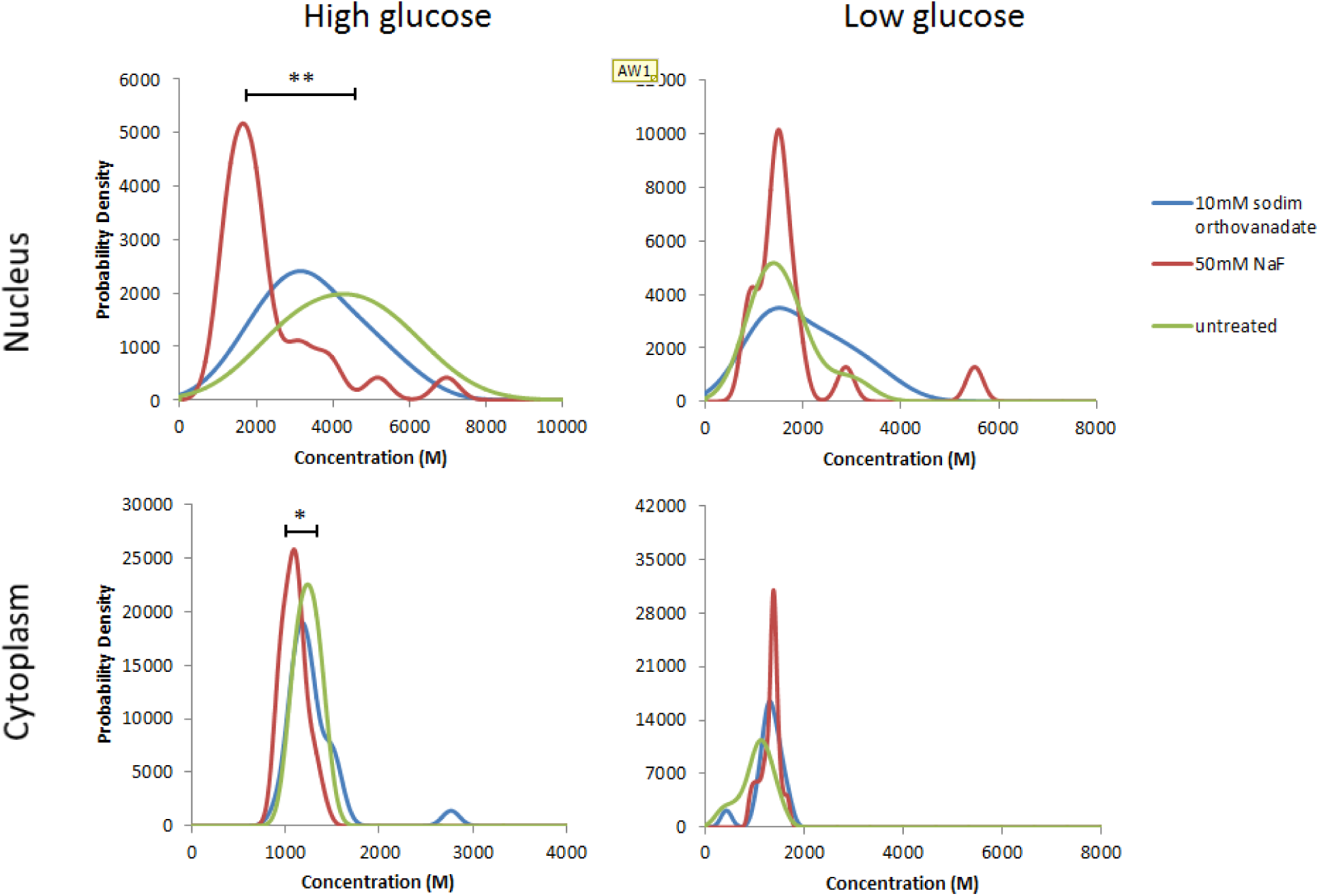
Inhibition of phosphatases reduces concentration of nuclear Mig1 in high glucose. Mig1 concentration in the nucleus and the cytoplasm under 4% (High) or 0.2% (Low) glucose conditions. Cultures were exposed to 10mM Na_3_VO_4_ or to 10mM NaF for 2h prior to imaging. Student t-test (p<0.05). *p>0.05, **p<0.001

Untreated cells contained similar concentrations of Mig1 in nucleus and cytoplasm as previously reported (Wollman and Leake 2015), with elevated concentration in the nucleus at high glucose. No significant effect of Na_3_VO_4_ on Mig1 nuclear and cytoplasmic concentration was identified at high or low glucose (Student’s t-test, P>0.05) (Figure 5), despite the disruption of wild type phosphorylation levels. However, incubation with 50mM NaF at high glucose resulted in heterogeneous behavior with many cells showing a significant reduction of Mig1 in the nucleus (Student’s t-test, P<0.001). A slight reduction was also observed in the cytoplasm (Student’s t-test, P<0.05). Decrease of the general protein level upon incubation with NaF is consistent with a role of fluoride ions in inhibition of protein synthesis initiation (Eckert 1975, O’Rourke and Godchaux 1975). As expected inhibition of Snf1 by 1NM-PP1 upon glucose limitation resulted in concentrated levels of Mig1 in the nucleus similar to high glucose conditions (Figure 4B, Supplementary Figure 3).

## Discussion

Mig1 has several phosphorylation sites mainly located on serine residues and a few on threonine (full list: http://yeastgenome.org/locus/S000003003/protein). Ser222 was proposed to be involved in cytoplasmic import of Mig1 upon glucose depletion (DeVit and Johnston 1999), Ser311 was shown to be required for *SUC2* derepression under glucose depletion and also for the interaction between Mig1 and Hxk2, which seems to be involved in glucose repression (Ahuatzi, et al. 2007). Our data support previous reports that Snf1-dependent phosphorylation is essential for Mig1 relocalization. Activated Snf1 mediates phosphorylation of Mig1. Here we show that inhibition of the Snf1 kinase activity results in Mig1 dephosphorylation and nuclear import.

A protein phosphatase 1, Glc7-Reg1, dephosphorylates Snf1 on Thr210 in high glucose conditions (Ludin, et al. 1998, McCartney and Schmidt 2001). In high glucose, the Reg1 subunit is dephosphorylated and associated with Glc7, which results in recruitment of the phosphatase activity towards Snf1 (Sanz, et al. 2000). It has been suggested that the Glc7-Reg1 phosphatase participates in Mig1 dephosphorylation under glucose repletion (McCartney and Schmidt 2001, Rubenstein, et al. 2008). Here we confirm that indeed Reg1 plays a role in Mig1 dephosphorylation and that this event is controlled by the presence of glucose. However, lack of Reg1 does not seem to affect the Mig1 phosphorylation state in high or low glucose at steady-state conditions. Neither does pharmacological inhibition of Snf1 result in fully dephosphorylated Mig1. Only the combination of Snf1 inhibition and rapid shift to high glucose result in a Reg1-dependent strong Mig1 dephosphorylation. Hence, it appears that there are two unrelated mechanisms of Mig1 dephosphorylation. A glucose-and Glc7-Reg1 independent mechanism that becomes apparent on Snf1 inhibition as well as a glucose-and Glc7-Reg1 dependent mechanism.

Vanadate ions from NaVO_4_ and Na_3_VO_4_ have been shown to mediate phosphorylation of tumor suppressor p53 protein (Suzuki, et al. 2007). Vanadate compounds were also reported to inhibit protein phosphatases other than tyrosine specific ones (Parra-Diaz, et al. 1995, Reiter, et al. 2002). Tyrosine de/phosphorylation is a rare event in yeast: there are no true protein tyrosine kinases and just a few phosphatases (Chi, et al. 2007). A typical artificial substrate for Tyr-specific kinases contains a poly-(Glu-Tyr) motif. A preferred substrate sequence for Tyr-specific kinases in eukaryotic cells has been reported to contain hydrophobic and acidic residues downstream and upstream of a tyrosine, respectively (Blom, et al. 1999, Miller 2003). Our results suggest that vanadate ions cause constant phosphorylation of Mig1 regardless of glucose presence but no difference in Mig1 localization compared to untreated cells. Mig1 has six tyrosine residues. One of them, Tyr358, shares some sequence similarity with reported human tyrosine phosphorylation sites (Supplementary Figure 4) identified by PhosphoMotif Finder Software (Amanchy, et al. 2007). However, the general amino acid composition of the motif does not look like a preferred substrate sequence for tyrosine phosphorylation.

Vanadate ions might potentially trigger Mig1 phosphorylation on Ser/Thr sites that are not responsible for Mig1 nuclear export. The opposite effect might be achieved by NaF where Mig1 phosphorylation status remains similar to untreated cells but Mig1 fails to be mainly located in the nucleus under glucose repletion. Retardation in protein migration upon SDS-Page separation cannot identify which sites are phosphorylated, thus, the exact effect of vanadate ions on Mig1phopshorylation remains to be determined.

NaF is an inhibitor of serine-threonine phosphatases (Shenolikar and Nairn 1991), although, it was reported that NaF does not completely inhibit general level of serine/threonine phosphatases activity. Thus, the exposure to this compound might result in phosphorylation of one or very few residues of Mig1, which is not captured by western blotting but sufficient to keep the protein in the cytoplasm. On the other hand, NaF might not affect Mig1 directly but mediates phosphorylation of another agent which in this state binds to Mig1 preventing its nuclear import. In fact, Hxk2 was suggested to be responsible for Mig1 relocalization to the nucleus in glucose repletion (Ahuatzi, et al. 2007). Hxk2 is a serine/threonine kinase which together with Hxk1 and Glk1 phosphorylate glucose and becomes phosphorylated itself in the presence of alternative carbon sources (Randez-Gil, et al. 1998). Therefore, NaF might act on Mig1 localization through phosphorylation of Hxk2. Further work is needed to confirm this.

Our results indicate that Mig1 dephosphorylation is triggered by a component dependent on tyrosine-specific dephosphorylation. Large scale analysis identified various phosphorylation sites of Reg1 mainly on serine and threonine residues but also on tyrosine 480 (Chi, et al. 2007). Although it has been proposed that Reg1 dephosphorylation is promoted by the serine/threonine phosphatase Glc7 (Sanz, et al. 2000), our data suggest that Tyr480 phosphorylation of Reg1 might be crucial for the PP1 action towards Mig1 dephosphorylation. However, there could be another tyrosine-phosphorylation dependent agent acting upstream of Mig1-dephosphorylating phosphatase.

Overall, Mig1 de/phosphorylation appears to be a complex process controlled in glucose-dependent and independent manner by the orchestrated action of protein phosphatases and protein kinases.

### Funding

This work was supported by European Commission via Marie Curie-Network for Initial training ISOLATE (Grant agreement nr: 289995), Swedish Research Council, the Biological Physical Sciences Institute, Royal Society, MRC (grant MR/K01580X/1) and the Royal Society Newton International Fellowship.

---

**Supplementary Figure 1.**
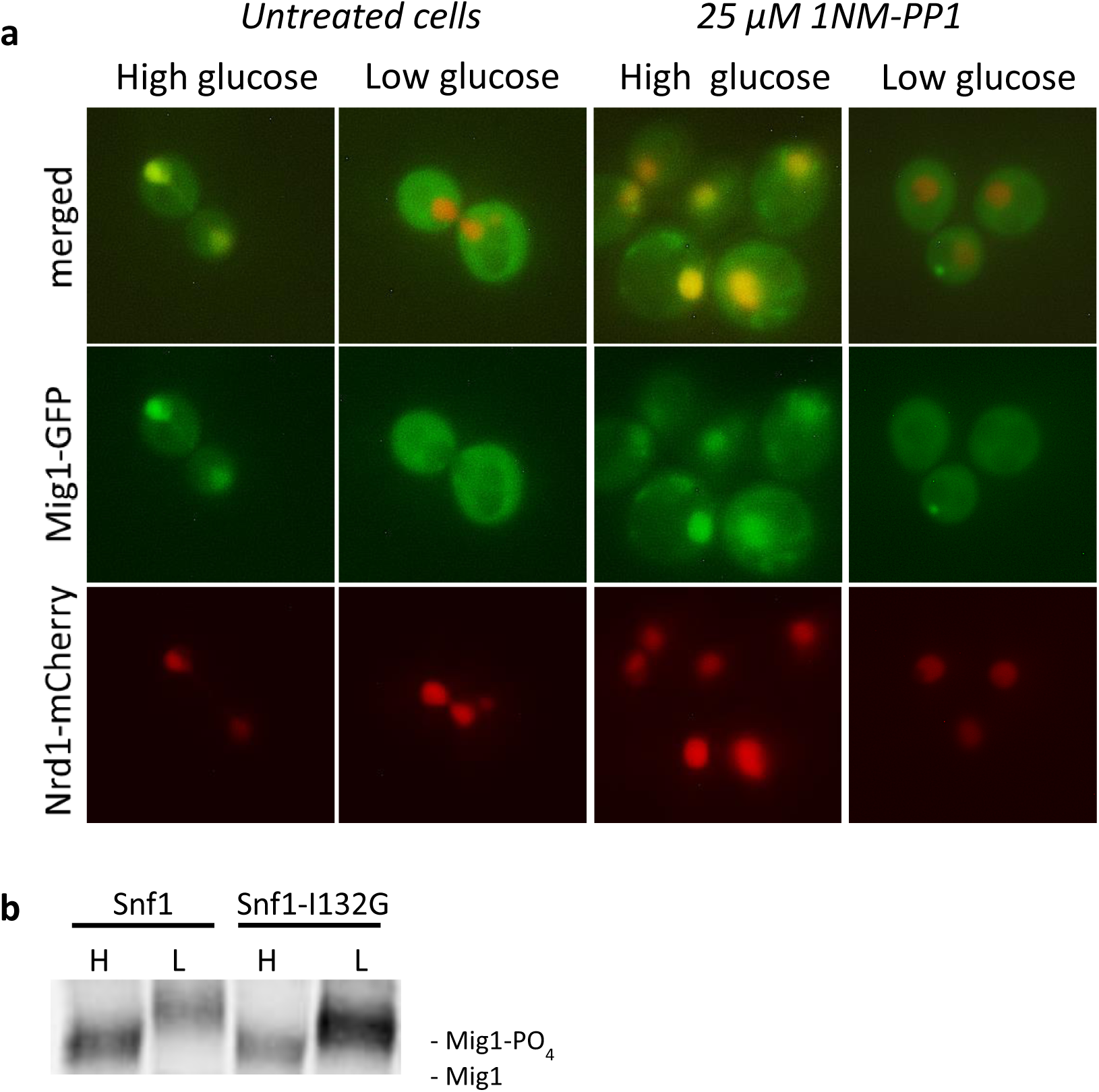
I132G mutation makes Snf1 sensitive to the inhibitor but does not affect its function. **(a)** Cells with the wild type form of Snf1 and expressing Mig1-GFP and Nrd1-mCherry as a marker for the nucleus were incubated in different glucose conditions and then exposed to 25 μM 1NM-PP1. **(b)** Phosphorylation status of Mig1 obtained from 70 μg of total protein extracts from cells expressing Mig1-HA and the wild type or analogue-sensitive version of Snf1, Snf1-I132G, under 4% (H) or 0.2% (L) glucose conditions.

**Supplementary Figure 2.**
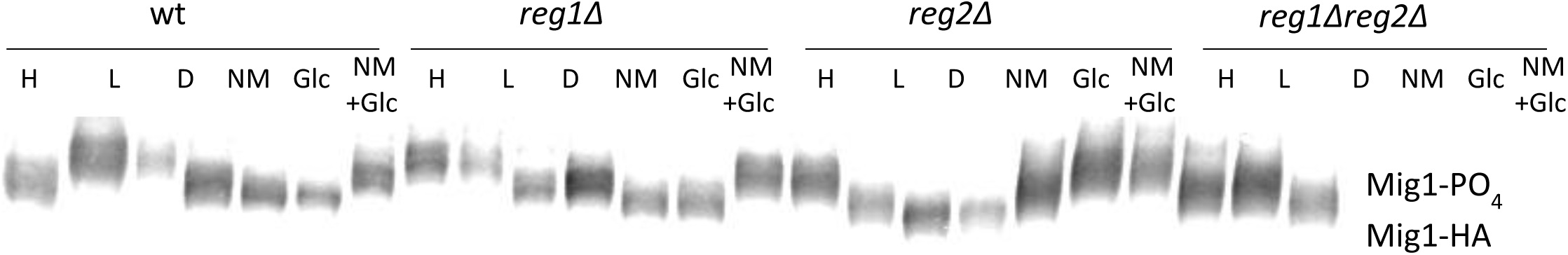
Mig1 dephosphorylation by Glc7-Reg1 is mediated by glucose. The phosphorylation status of Mig1 was determined by Western blotting of protein extracts from cells expressing HA-tagged Mig1 and an ATP-analogue sensitive Snf1, Snf1-I132G, grown in 4% glucose, shifted to 4% (H) and 0.2% glucose (L) for 1h. 0.2% glucose cells were subsequently treated with DMSO (D), the inhibitor of Snf1-I132G, 25uM 1NM-PP1 (NM), 4% glucose (Glc) for 5 min or 4% glucose after 1NM-PP1 treatment (NM+Glc).

**Supplementary Figure 3.**
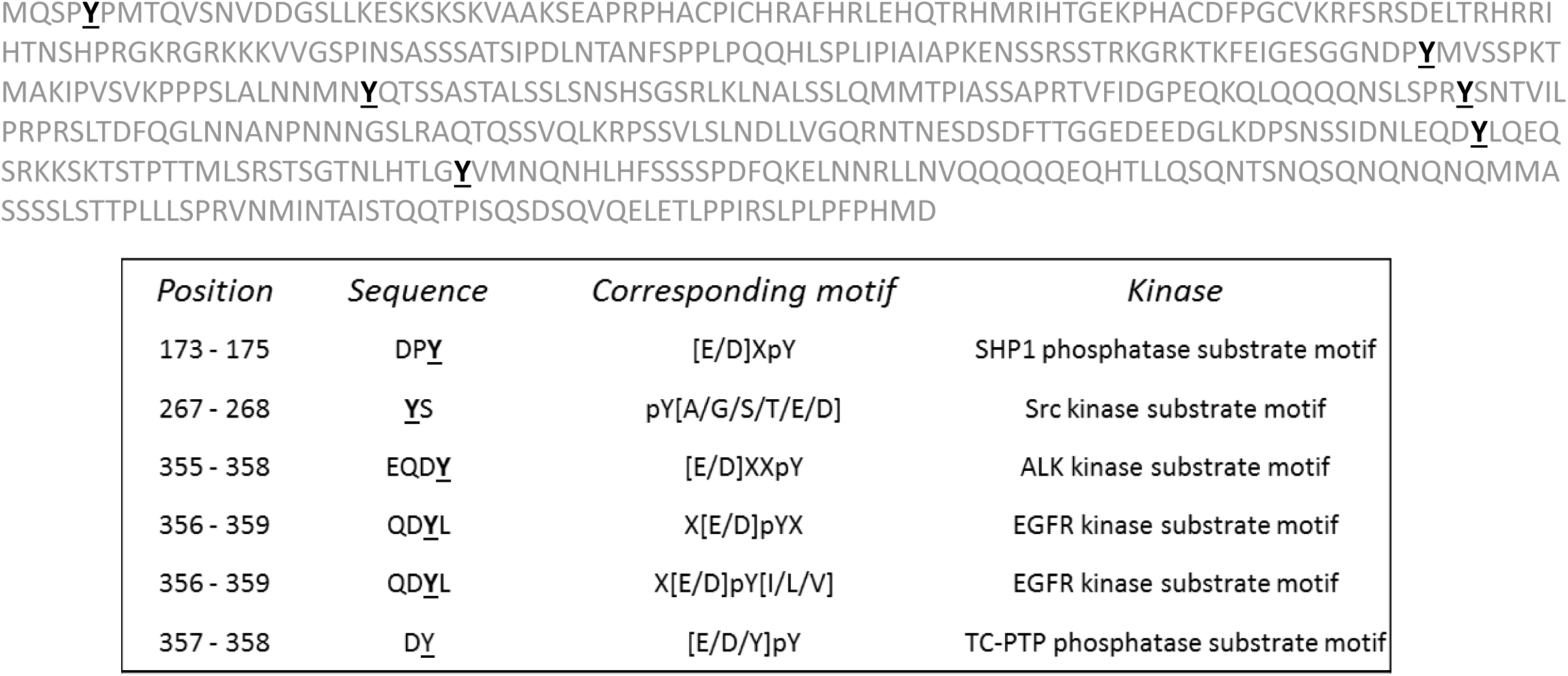
Tyrosine residues in Mig1 sequence and their similarity to reported mammalian tyrosine kinase/phosphatase motifs.

## References

Ahuatzi, D., A. Riera, R. Pelaez, P. Herrero, and F. Moreno. Hxk2 regulates the phosphorylation state of Mig1 and therefore its nucleocytoplasmic distribution. J Biol Chem 2007; vol. 282, no. 7, pp. 4485–93.

Amanchy, R., B. Periaswamy, S. Mathivanan, R. Reddy, S. G. Tattikota, and A. Pandey. A curated compendium of phosphorylation motifs. Nat Biotechnol 2007; vol. 25, no. 3, pp. 285–6.

Beattie, T. R., N. Kapadia, E. Nicolas, S. Uphoff, A. J. Wollman, M. C. Leake, and R. Reyes-Lamothe. Frequent exchange of the DNA polymerase during bacterial chromosome replication. Elife 2017; vol. 6.

Bendrioua, L., M. Smedh, J. Almquist, M. Cvijovic, M. Jirstrand, M. Goksor, C. B. Adiels, and S. Hohmann. Yeast AMP-activated protein kinase monitors glucose concentration changes and absolute glucose levels. J Biol Chem 2014; vol. 289, no. 18, pp. 12863–75.

Blom, N., S. Gammeltoft, and S. Brunak. Sequence and structure-based prediction of eukaryotic protein phosphorylation sites. J Mol Biol 1999; vol. 294, no. 5, pp. 1351–62.

Carling, D., K. Aguan, A. Woods, A. J. Verhoeven, R. K. Beri, C. H. Brennan, C. Sidebottom, M. D. Davison, and J. Scott. Mammalian AMP-activated protein kinase is homologous to yeast and plant protein kinases involved in the regulation of carbon metabolism. J Biol Chem 1994; vol. 269, no. 15, pp. 11442–8.

Carlson, M. Glucose repression in yeast. Curr Opin Microbiol 1999; vol. 2, no. 2, pp. 202–7.

Carlson, M., B. C. Osmond, and D. Botstein. Mutants of yeast defective in sucrose utilization. Genetics 1981; vol. 98, no. 1, pp. 25–40.

Celenza, J. L., and M. Carlson. A yeast gene that is essential for release from glucose repression encodes a protein kinase. Science 1986; vol. 233, no. 4769, pp. 1175–80.

Chi, A., C. Huttenhower, L. Y. Geer, J. J. Coon, J. E. Syka, D. L. Bai, J. Shabanowitz, D. J. Burke, O. G. Troyanskaya, and D. F. Hunt. Analysis of phosphorylation sites on proteins from Saccharomyces cerevisiae by electron transfer dissociation (ETD) mass spectrometry. Proc Natl Acad Sci U S A 2007; vol. 104, no. 7, pp. 2193–8.

Crans, D. C., J. J. Smee, E. Gaidamauskas, and L. Yang. The chemistry and biochemistry of vanadium and the biological activities exerted by vanadium compounds. Chem Rev 2004; vol. 104, no. 2, pp. 849–902.

De Vit, M. J., J. A. Waddle, and M. Johnston. Regulated nuclear translocation of the Mig1 glucose repressor. Mol Biol Cell 1997; vol. 8, no. 8, pp. 1603–18.

DeVit, M. J., and M. Johnston. The nuclear exportin Msn5 is required for nuclear export of the Mig1 glucose repressor of Saccharomyces cerevisiae. Curr Biol 1999; vol. 9, no. 21, pp. 1231–41.

Eckert, J. Diffuse hair loss and psychiatric disturbance. Acta Derm Venereol 1975; vol. 55, no. 2, pp. 147–9.

Elbing, K., R. R. McCartney, and M. C. Schmidt. Purification and characterization of the three Snf1-activating kinases of Saccharomyces cerevisiae. Biochem J 2006; vol. 393, no. Pt 3, pp. 797–805.

Frederick, D. L., and K. Tatchell. The REG2 gene of Saccharomyces cerevisiae encodes a type 1 protein phosphatase-binding protein that functions with Reg1p and the Snf1 protein kinase to regulate growth. Mol Cell Biol 1996; vol. 16, no. 6, pp. 2922–31.

Gietz, R. D., and R. H. Schiestl. Frozen competent yeast cells that can be transformed with high efficiency using the LiAc/SS carrier DNA/PEG method. Nat Protoc 2007; vol. 2, no. 1, pp. 1–4.

Gordon, J. A. Use of vanadate as protein-phosphotyrosine phosphatase inhibitor. Methods Enzymol 1991; vol. 201, pp. 477–82.

Hong, S. P., F. C. Leiper, A. Woods, D. Carling, and M. Carlson. Activation of yeast Snf1 and mammalian AMP-activated protein kinase by upstream kinases. Proc Natl Acad Sci U S A 2003; vol. 100, no. 15, pp. 8839–43.

Hu, Z., J. O. Nehlin, H. Ronne, and C. A. Michels. MIG1-dependent and MIG1-independent glucose regulation of MAL gene expression in Saccharomyces cerevisiae. Curr Genet 1995; vol. 28, no. 3, pp. 258–66.

Jiang, H., K. Tatchell, S. Liu, and C. A. Michels. Protein phosphatase type-1 regulatory subunits Reg1p and Reg2p act as signal transducers in the glucose-induced inactivation of maltose permease in Saccharomyces cerevisiae. Mol Gen Genet 2000; vol. 263, no. 3, pp. 411–22.

Johnston, M., J. S. Flick, and T. Pexton. Multiple mechanisms provide rapid and stringent glucose repression of GAL gene expression in Saccharomyces cerevisiae. Mol Cell Biol 1994; vol. 14, no. 6, pp. 3834–41.

Knight, Z. A., and K. M. Shokat. Chemical genetics: where genetics and pharmacology meet. Cell 2007; vol. 128, no. 3, pp. 425–30.

Korbecki, J., I. Baranowska-Bosiacka, I. Gutowska, and D. Chlubek. Biochemical and medical importance of vanadium compounds. Acta Biochim Pol 2012; vol. 59, no. 2, pp. 195–200.

Leake, M. C., J. H. Chandler, G. H. Wadhams, F. Bai, R. M. Berry, and J. P. Armitage. Stoichiometry and turnover in single, functioning membrane protein complexes. Nature 2006; vol. 443, no. 7109, pp. 355–8.

Ludin, K., R. Jiang, and M. Carlson. Glucose-regulated interaction of a regulatory subunit of protein phosphatase 1 with the Snf1 protein kinase in Saccharomyces cerevisiae. Proc Natl Acad Sci U S A 1998; vol. 95, no. 11, pp. 6245–50.

Maziarz, M., A. Shevade, L. Barrett, and S. Kuchin. Springing into Action: Reg2 Negatively Regulates Snf1 Protein Kinase and Facilitates Recovery from Prolonged Glucose Starvation in Saccharomyces cerevisiae. Appl Environ Microbiol 2016; vol. 82, no. 13, pp. 3875–85.

McCartney, R. R., and M. C. Schmidt. Regulation of Snf1 kinase. Activation requires phosphorylation of threonine 210 by an upstream kinase as well as a distinct step mediated by the Snf4 subunit. J Biol Chem 2001; vol. 276, no. 39, pp. 36460–6.

Miller, H., Z. Zhou, A. J. Wollman, and M. C. Leake. Superresolution imaging of single DNA molecules using stochastic photoblinking of minor groove and intercalating dyes. Methods 2015; vol. 88, pp. 81–8.

Miller, W. T. Determinants of substrate recognition in nonreceptor tyrosine kinases. Acc Chem Res 2003; vol. 36, no. 6, pp. 393–400.

Nath, N., R. R. McCartney, and M. C. Schmidt. Yeast Pak1 kinase associates with and activates Snf1. Mol Cell Biol 2003; vol. 23, no. 11, pp. 3909–17.

Needham, P. G., and R. J. Trumbly. In vitro characterization of the Mig1 repressor from Saccharomyces cerevisiae reveals evidence for monomeric and higher molecular weight forms. Yeast 2006; vol. 23, no. 16, pp. 1151–66.

Nehlin, J. O., M. Carlberg, and H. Ronne. Control of yeast GAL genes by MIG1 repressor: a transcriptional cascade in the glucose response. EMBO J 1991; vol. 10, no. 11, pp. 3373–7.

O’Rourke, J. C., and W. Godchaux, 3rd. Fluoride inhibition of the initiation of protein synthesis in the reticulocyte lysate cell-free system. J Biol Chem 1975; vol. 250, no. 9, pp. 3443–50.

Papamichos-Chronakis, M., T. Gligoris, and D. Tzamarias. The Snf1 kinase controls glucose repression in yeast by modulating interactions between the Mig1 repressor and the Cyc8-Tup1 co-repressor. EMBO Rep 2004; vol. 5, no. 4, pp. 368–72.

Parra-Diaz, D., Q. Wei, E. Y. Lee, L. Echegoyen, and D. Puett. Binding of vanadium (IV) to the phosphatase calcineurin. FEBS Lett 1995; vol. 376, no. 1-2, pp. 58–60.

Plank, M., G. H. Wadhams, and M. C. Leake. Millisecond timescale slimfield imaging and automated quantification of single fluorescent protein molecules for use in probing complex biological processes. Integr Biol (Camb) 2009; vol. 1, no. 10, pp. 602–12.

Randez-Gil, F., P. Sanz, K. D. Entian, and J. A. Prieto. Carbon source-dependent phosphorylation of hexokinase PII and its role in the glucose-signaling response in yeast. Mol Cell Biol 1998; vol. 18, no. 5, pp. 2940–8.

Reiter, N. J., D. J. White, and F. Rusnak. Inhibition of bacteriophage lambda protein phosphatase by organic and oxoanion inhibitors. Biochemistry 2002; vol. 41, no. 3, pp. 1051–9.

Reyes-Lamothe, R., D. J. Sherratt, and M. C. Leake. Stoichiometry and architecture of active DNA replication machinery in Escherichia coli. Science 2010; vol. 328, no. 5977, pp. 498–501.

Rubenstein, E. M., R. R. McCartney, C. Zhang, K. M. Shokat, M. K. Shirra, K. M. Arndt, and M. C. Schmidt. Access denied: Snf1 activation loop phosphorylation is controlled by availability of the phosphorylated threonine 210 to the PP1 phosphatase. J Biol Chem 2008; vol. 283, no. 1, pp. 222–30.

Ruiz, A., Y. Liu, X. Xu, and M. Carlson. Heterotrimer-independent regulation of activation-loop phosphorylation of Snf1 protein kinase involves two protein phosphatases. Proc Natl Acad Sci U S A 2012; vol. 109, no. 22, pp. 8652–7.

Sanz, P., G. R. Alms, T. A. Haystead, and M. Carlson. Regulatory interactions between the Reg1-Glc7 protein phosphatase and the Snf1 protein kinase. Mol Cell Biol 2000; vol. 20, no. 4, pp. 1321–8.

Shashkova, S., N. Welkenhuysen, and S. Hohmann. Molecular communication: crosstalk between the Snf1 and other signaling pathways. FEMS Yeast Res 2015; vol. 15, no. 4, p. fov026.

Shenolikar, S., and A. C. Nairn. Protein phosphatases: recent progress. Adv Second Messenger Phosphoprotein Res 1991; vol. 23, pp. 1–121.

Shirra, M. K., R. R. McCartney, C. Zhang, K. M. Shokat, M. C. Schmidt, and K. M. Arndt. A chemical genomics study identifies Snf1 as a repressor of GCN4 translation. J Biol Chem 2008; vol. 283, no. 51, pp. 35889–98.

Suzuki, K., K. Inageda, G. Nishitai, and M. Matsuoka. Phosphorylation of p53 at serine 15 in A549 pulmonary epithelial cells exposed to vanadate: involvement of ATM pathway. Toxicol Appl Pharmacol 2007; vol. 220, no. 1, pp. 83–91.

Treitel, M. A., and M. Carlson. Repression by SSN6-TUP1 is directed by MIG1, a repressor/activator protein. Proc Natl Acad Sci U S A 1995; vol. 92, no. 8, pp. 3132–6.

Treitel, M. A., S. Kuchin, and M. Carlson. Snf1 protein kinase regulates phosphorylation of the Mig1 repressor in Saccharomyces cerevisiae. Mol Cell Biol 1998; vol. 18, no. 11, pp. 6273–80.

Tu, J., and M. Carlson. The GLC7 type 1 protein phosphatase is required for glucose repression in Saccharomyces cerevisiae. Mol Cell Biol 1994; vol. 14, no. 10, pp. 6789–96.

Wang, P., A. D. Verin, A. Birukova, L. I. Gilbert-McClain, K. Jacobs, and J. G. Garcia. Mechanisms of sodium fluoride-induced endothelial cell barrier dysfunction: role of MLC phosphorylation. Am J Physiol Lung Cell Mol Physiol 2001; vol. 281, no. 6, pp. L1472–83.

Wollman, A. J., and M. C. Leake. Millisecond single-molecule localization microscopy combined with convolution analysis and automated image segmentation to determine protein concentrations in complexly structured, functional cells, one cell at a time. Faraday Discuss 2015; vol. 184, pp. 401–24.

Wollman, A. J., A. H. Syeda, P. McGlynn, and M. C. Leake. Single-Molecule Observation of DNA Replication Repair Pathways in E. coli. Adv Exp Med Biol 2016; vol. 915, pp. 5–16.

Wollman, Adam J.M., Leake, Mark C., ‘Single-Molecule Narrow-Field Microscopy of Protein–DNA Binding Dynamics in Glucose Signal Transduction of Live Yeast Cells’, Chromosome Architecture Methods and Protocols. Methods in Molecular Biology, 2016, 2016, pp. X, 290 p. 77 illus., 65 illus. in color.

Wu, J., and R. J. Trumbly. Multiple regulatory proteins mediate repression and activation by interaction with the yeast Mig1 binding site. Yeast 1998; vol. 14, no. 11, pp. 985–1000.

Zaman, S., S. I. Lippman, L. Schneper, N. Slonim, and J. R. Broach. Glucose regulates transcription in yeast through a network of signaling pathways. Mol Syst Biol 2009; vol. 5, p. 245.

